## Supplementary data for "Deciphering ion transport and ATPase coupling in the intersubunit tunnel of KdpFABC"

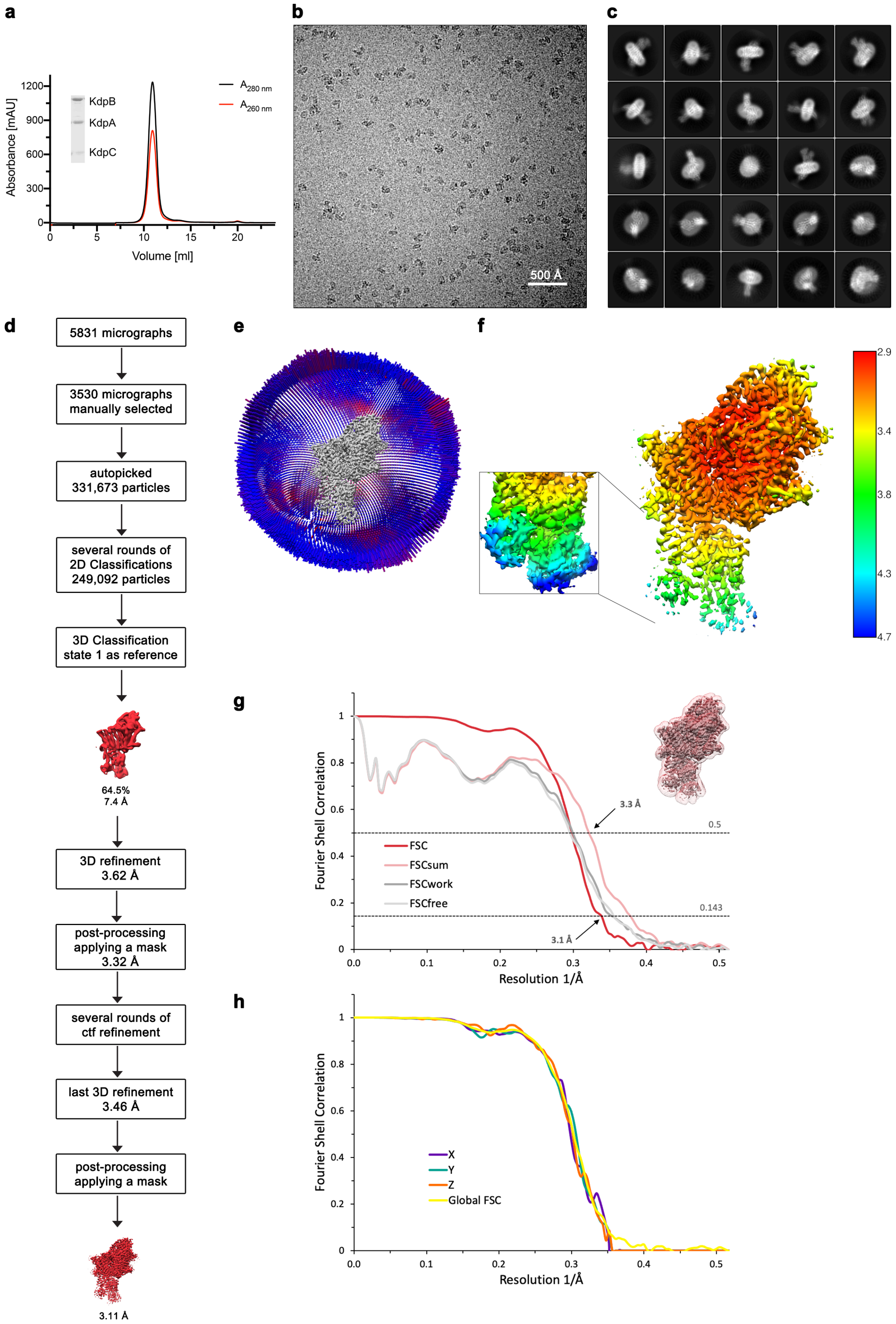

**Supplementary Figure 1: Cryo-EM analysis of $K^+$-loaded KdpFAB$_{D307N}$C in the E1·ATP state.** **a,** Purification of KdpFAB$_{D307N}$C. A representative SEC elution profile (Superdex 200 incr. 10/300) and SDS-PAGE are shown. **b,** Representative micrograph of the recorded data. **c,** 2D class averages of vitrified KdpFAB$_{D307N}$C in the presence of 50 mM KCl, and 5 mM AMPPCP. **d,** Image processing workflow as described in the methods section. **e,** Angular distribution plot of particles included in the unsymmetrized 3D reconstruction for KdpFABC. The number of particles with the respective orientation is represented by length and color of the cylinders (long and red – high number of particles; short and blue – low number of particles). **f,** Final reconstruction map colored by local resolution as estimated by RELION[52]. **g,** FSC plot used for resolution estimation and model validation. The gold-standard FSC plot between two separately refined half-maps is shown in red and indicates a final resolution of 3.1 Å. The FSC model validation curves for FSCsum, FSCwork and FSCfree, as described in the methods, are shown in light red, dark grey and light grey respectively. A thumbnail of the mask used for FSC calculation overlaid on the map is shown in the upper right corner. Dashed lines indicate the FSC thresholds used for FSC (0.143) and for FSCsum (0.5). **h,** Anisotropy estimation plot of the final map. The global FSC curve is represented in yellow. The directional FSCs along the x-, y- and z-axes are displayed in blue, green and red, respectively.

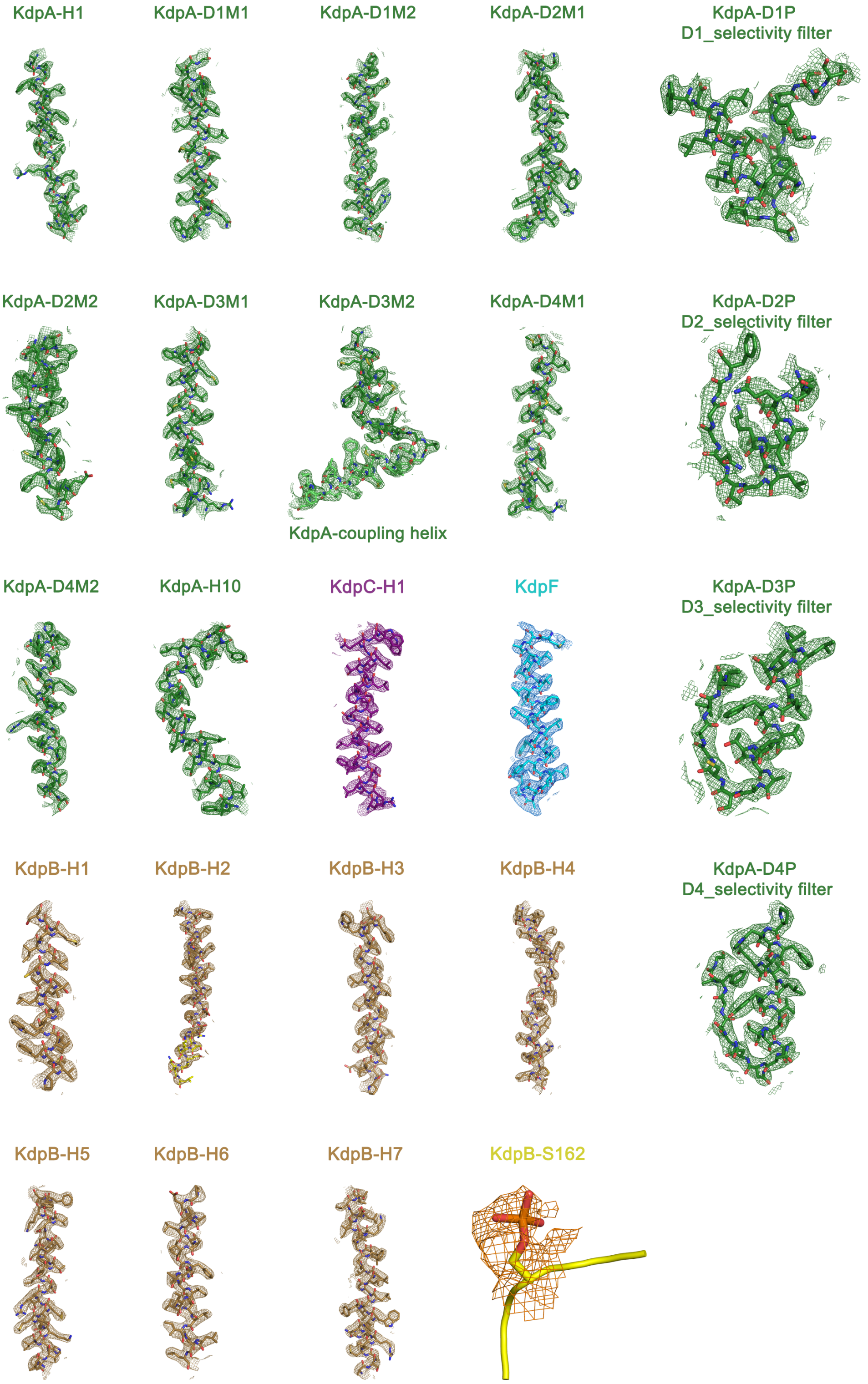

**Supplementary Figure 2: Cryo-EM densities of the membrane-inserted moieties and mutated residues of KdpFAB$_{D307N}$C.** TM helices 1-10 and selectivity filter pore loops of KdpA (green), TM helices 1-7 of KdpB (sand), the phosphorylated KdpB$_{S162}$ (yellow), TM helix of KdpC (purple), and KdpF (cyan) are fitted to the corresponding maps shown at 7σ.

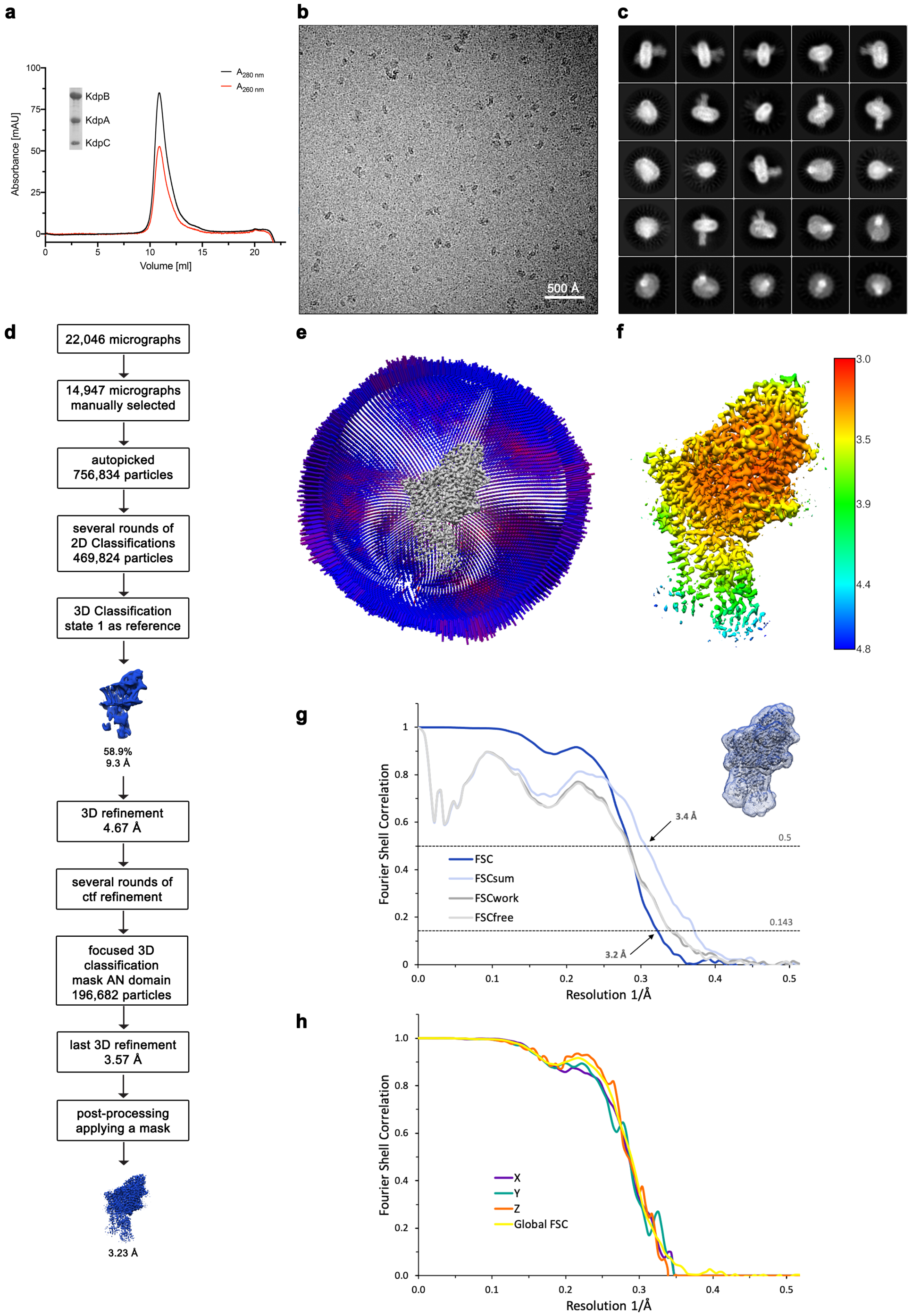

**Supplementary Figure 3: Cryo-EM analysis of Rb$^+$-loaded KdpFA$_{G232D}$B$_{S162A}$C in the E1·ATP state. a,** Purification of KdpFA$_{G232D}$B$_{S162A}$C. A representative SEC elution profile (Superdex 200 incr. 10/300) and SDS-PAGE are shown. **b,** Representative micrograph of the recorded data. **c,** 2D class averages of vitrified KdpFA$_{G232D}$B$_{S162A}$C in the presence of 1 mM ATP, 100 mM RbCl, and 10 mM AMPPCP. **d,** Image processing workflow as described in the methods section. **e,** Angular distribution plot of particles included in the unsymmetrized 3D reconstruction for KdpFABC. The number of particles with the respective orientation is represented by length and color of the cylinders (long and red – high number of particles; short and blue – low number of particles). **f,** Final reconstruction map colored by local resolution as estimated by RELION[52]. Notably, the local resolution in the A domain is lower than in the structure of $KdpFAB_{D307N}C$, reflecting effects observed in other structures containing mutation $KdpB_{S162A}$. This suggests that phosphorylation of this residue stabilizes the conformation of the A domain. **g,** FSC plot used for resolution estimation and model validation. The gold-standard FSC plot between two separately refined half-maps is shown in blue and indicates a final resolution of 3.2 Å. The FSC model validation curves for FSCsum, FSCwork and FSCfree, as described in the methods, are shown in light blue, dark grey and light grey respectively. A thumbnail of the mask used for FSC calculation overlaid on the map is shown in the upper right corner. Dashed lines indicate the FSC thresholds used for FSC (0.143) and for FSCsum (0.5). **h,** Anisotropy estimation plot of the final map. The global FSC curve is represented in yellow. The directional FSCs along the x-, y- and z-axes are displayed in blue, green and red, respectively.

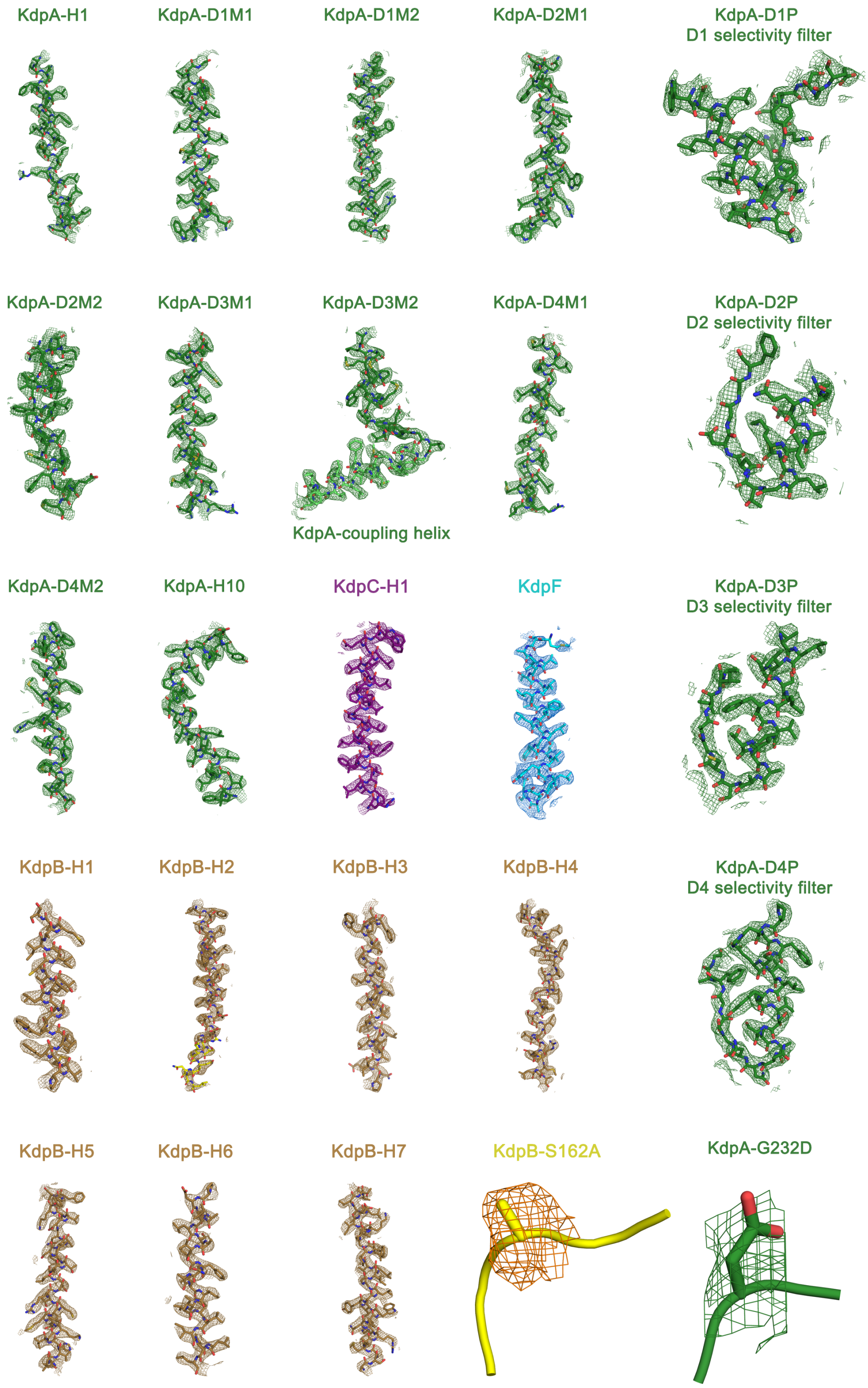

**Supplementary Figure 4: Cryo-EM densities of the membrane-inserted moieties and mutated residues of KdpFA$_{G232D}$B$_{S162A}$C.** TM helices 1-10 and selectivity filter pore loops of KdpA (green), the mutated SF residue KdpA$_{G232D}$, TM helices 1-7 of KdpB (sand), the phosphorylation-free KdpB$_{S162A}$ (yellow), TM helix of KdpC (purple), and KdpF (cyan) are fitted to the corresponding maps shown at 7σ.

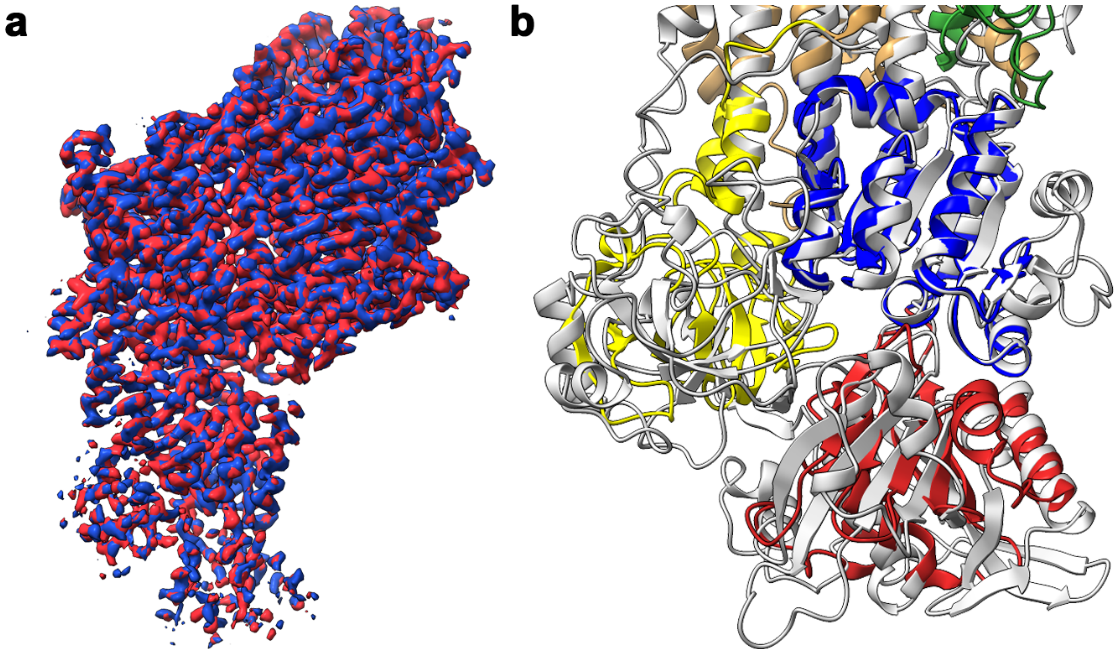

**Supplementary Figure 5: Conformational assignment of the presented KdpFABC structures.** **a,** Overlay of the cryo-EM maps generated for KdpFAB$_{D307N}$C (red) and KdpFA$_{G232D}$B$_{S162A}$C (blue), indicating an identical conformation in all regions of the complex. **b,** Overlay of the cytosolic domains from the K$^{+}$-loaded structure of KdpFAB$_{D307N}$C with AMPPCP and SERCA in an E1·ATP conformation [4XOU] (gray), verifying the assignment of the E1·ATP conformation to both structures presented here.

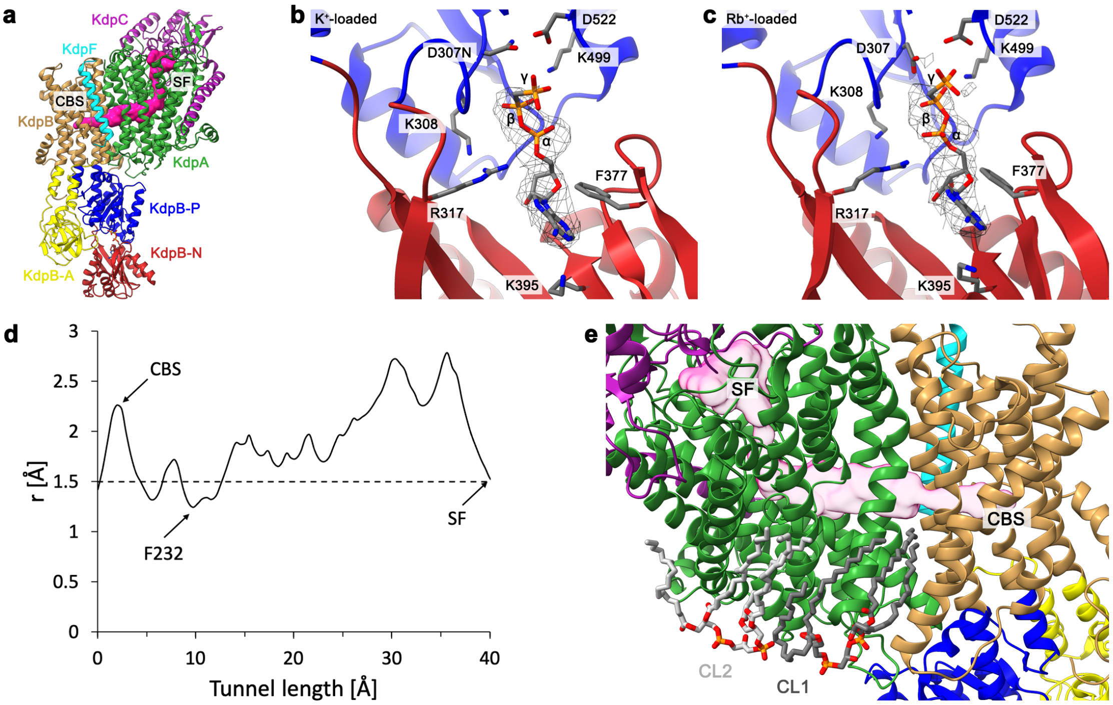

**Supplementary Figure 6: Structural features in the E1·ATP state. a,** Structural model of KdpFA$_{G232D}$B$_{S162A}$C in the ribbon representation. The KdpA pore entrance at the SF and intersubunit tunnel leading to the CBS are visualized as a pink surface. **b** and **c,** Nucleotide analog AMPPCP in the $K^+$- and $Rb^+$-loaded sample, respectively, modeled into its density (mesh), coordinated between the N and P domains, as was previously observed in other E1·ATP structures[11]. **d,** Radius of the intersubunit tunnel in $Rb^+$-loaded KdpFA$_{G232D}$B$_{S162A}$C. Like in the $K^+$-loaded sample, the tunnel is wide enough to permit $K^+$ (r=1.4 Å, dashed line), with a constriction at the KdpA SF and at KdpB$_{F232}$. **e,** Cardiolipin molecules CL1 and CL2 (dark and light grey, respectively) modeled in the cryo-EM structure of KdpFA$_{G232D}$B$_{S162A}$C.

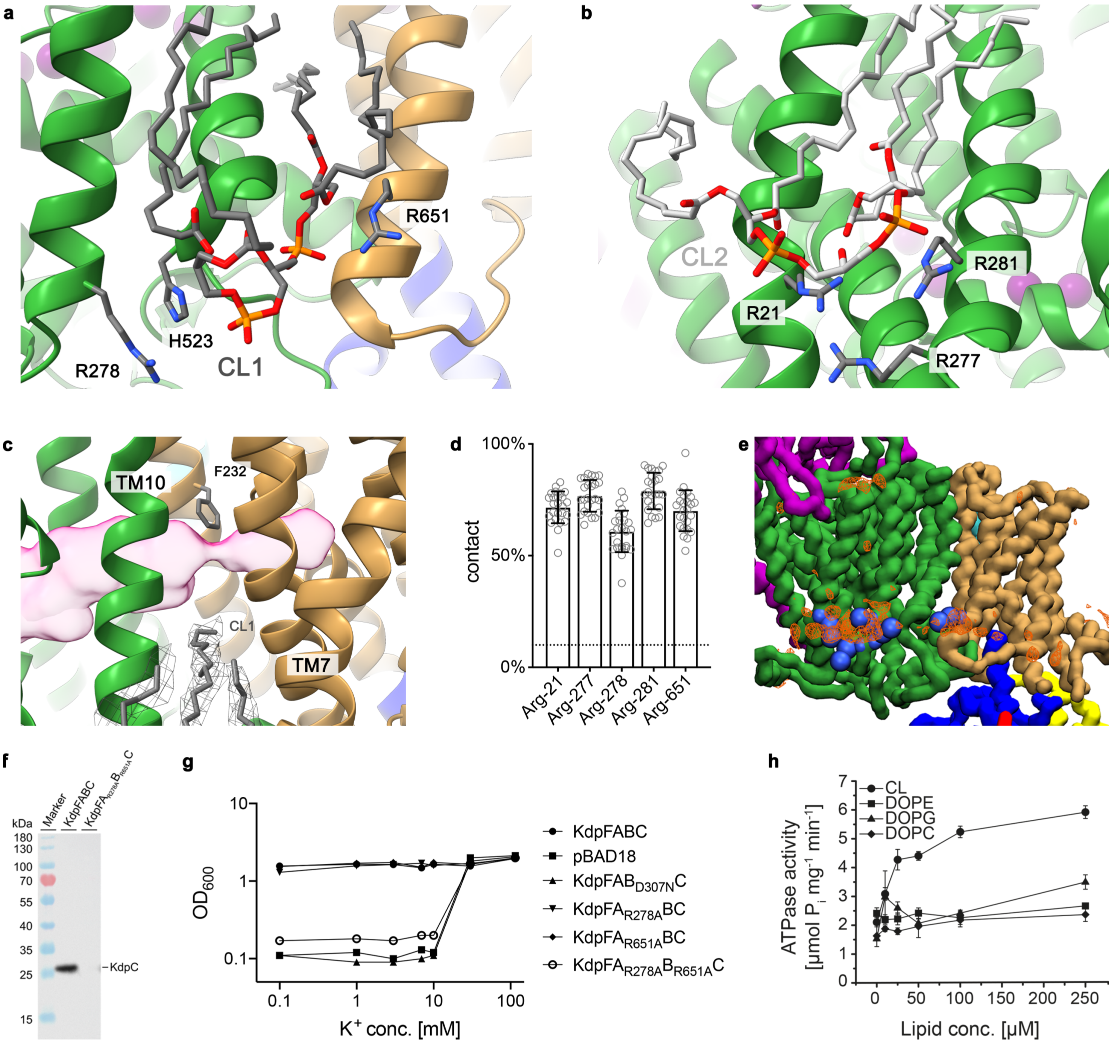

**Supplementary Figure 7: KdpFABC and cardiolipin. a,** The headgroup of CL 1 is coordinated by $KdpA_{R278}$, $KdpA_{H523}$, and $KdpB_{R651}$. **b,** The headgroup of CL2 is coordinated by $KdpA_{R21}$, $KdpA_{R277}$, and $KdpA_{R281}$. **c,** The hydrocarbon tail of CL1 that extends into the complex at the KdpA/KdpB interface lies below the path of the intersubunit tunnel at the site of constriction by $KdpB_{F232}$. **d,** Pooled CL contact probabilities of selected residues, from five representative structures in various E1 and E2 conformations ([5MRW], [6HRA], [6HRB], [7BGY], and [7NNL]), measured as proportion of frames the CL and respective residue are in contact, i.e. within a 0.6 nm cut-off distance. The concentration of CL in the membrane (10%) is shown as dotted line, indicating a significant accumulation of CL interactions at these residues. Error bars indicate the standard deviation from 25 independent simulations (5 each per structure). **e,** Identified arginine residues (blue spheres) with high CL binding propensity, as per panel c. The computed density for the CL molecules from the simulation data is shown as orange mesh. **f,** Anti-His Western Blot of whole-cell samples from the complementation set to an $OD_{600}$ of 10, detecting the His-tagged KdpC subunit. The CL binding site knockout variant is not expressed, showing the structural importance of CL binding for complex stability. **g,** Growth complementation assays for the analysis of the dependence of KdpFABC on cardiolipin. Growth measured by $OD_{600}$ after 24 h. Residues $KdpA_{R278}$ and $KdpB_{R651}$, which were identified by MD analysis as CL coordinating residues, were mutated to alanine to impair CL binding at the KdpA/KdpB interface. This double variant was unable to complement *E. coli* LB2003 cell growth under $K^+$ limitation, indicating a lack of $K^+$ translocation by the complex. In contrast, mutating the individual residues showed no effect. **h,** CL-dependent ATPase activity of KdpFABC. Rising CL conc. (0-250 μM) stimulate ATPase activity up to two-fold, while additions of PE, PG and PC do not change the ATPase activity of KdpFABC. Increased ATPase activity indicates an acceleration of the rate-limiting E1/E2 transition.

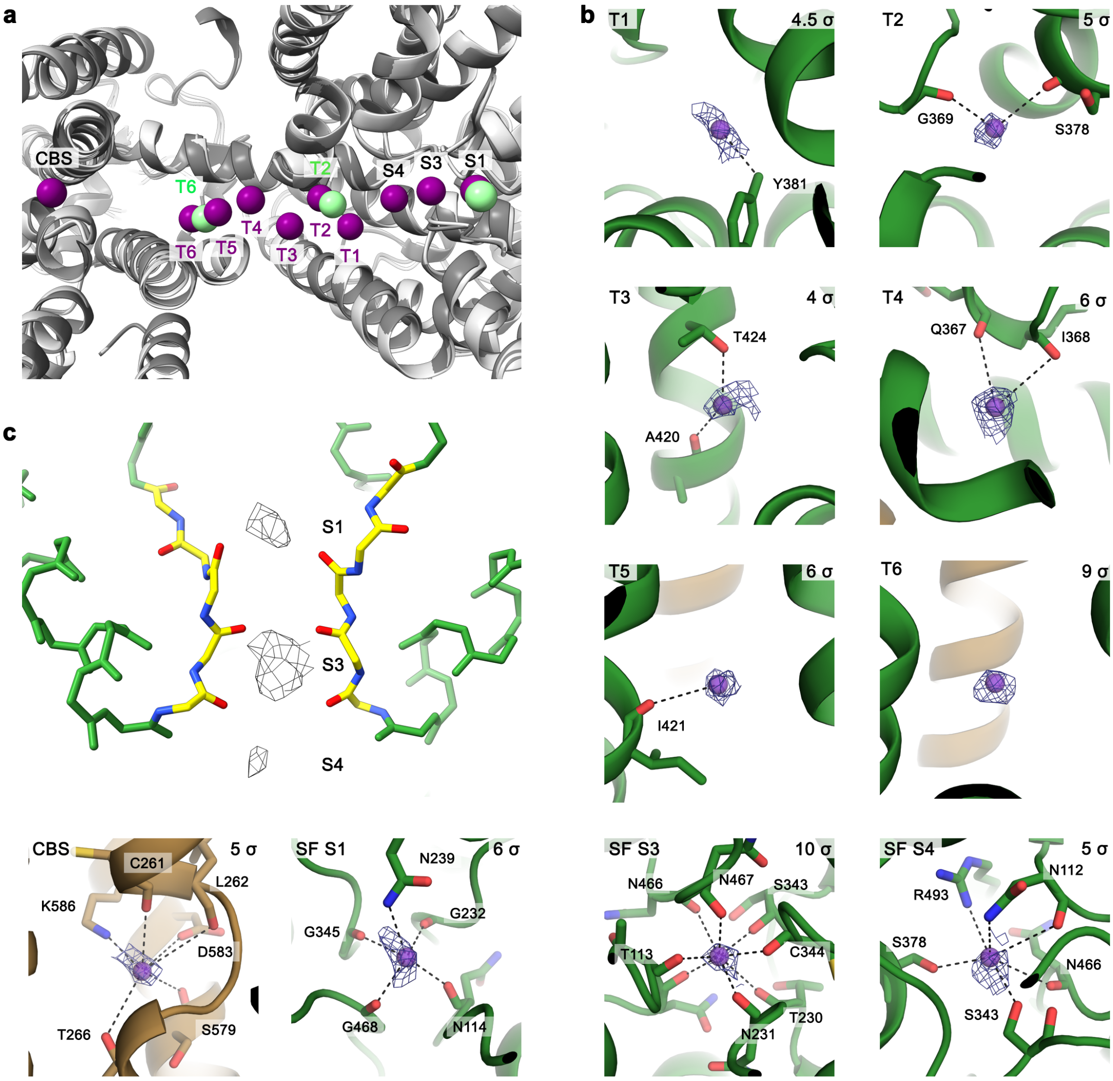

**Supplementary Figure 8: Ion coordination in $K^+$-loaded KdpFAB$_{D307N}$C. a,** Comparison of ion positions in the E1 structure [6HRA] and the $K^+$-loaded E1·ATP structure. $K^+$ ions in [6HRA] shown in green, newly identified $K^+$ ions in purple. The increased resolution and $K^+$ concentrations likely allow for the identification of additional densities. **b,** Coordination of $K^+$ ions in the intersubunit tunnel. No coordinating residues were identified for T6. Cryo-EM densities (mesh) shown with contour levels indicated in top right corner of each panel. **c,** SF of $K^+$-loaded KdpFAB$_{D307N}$C. Ion densities (mesh) in the S1, S3 and S4 sites are shown at 6 σ.

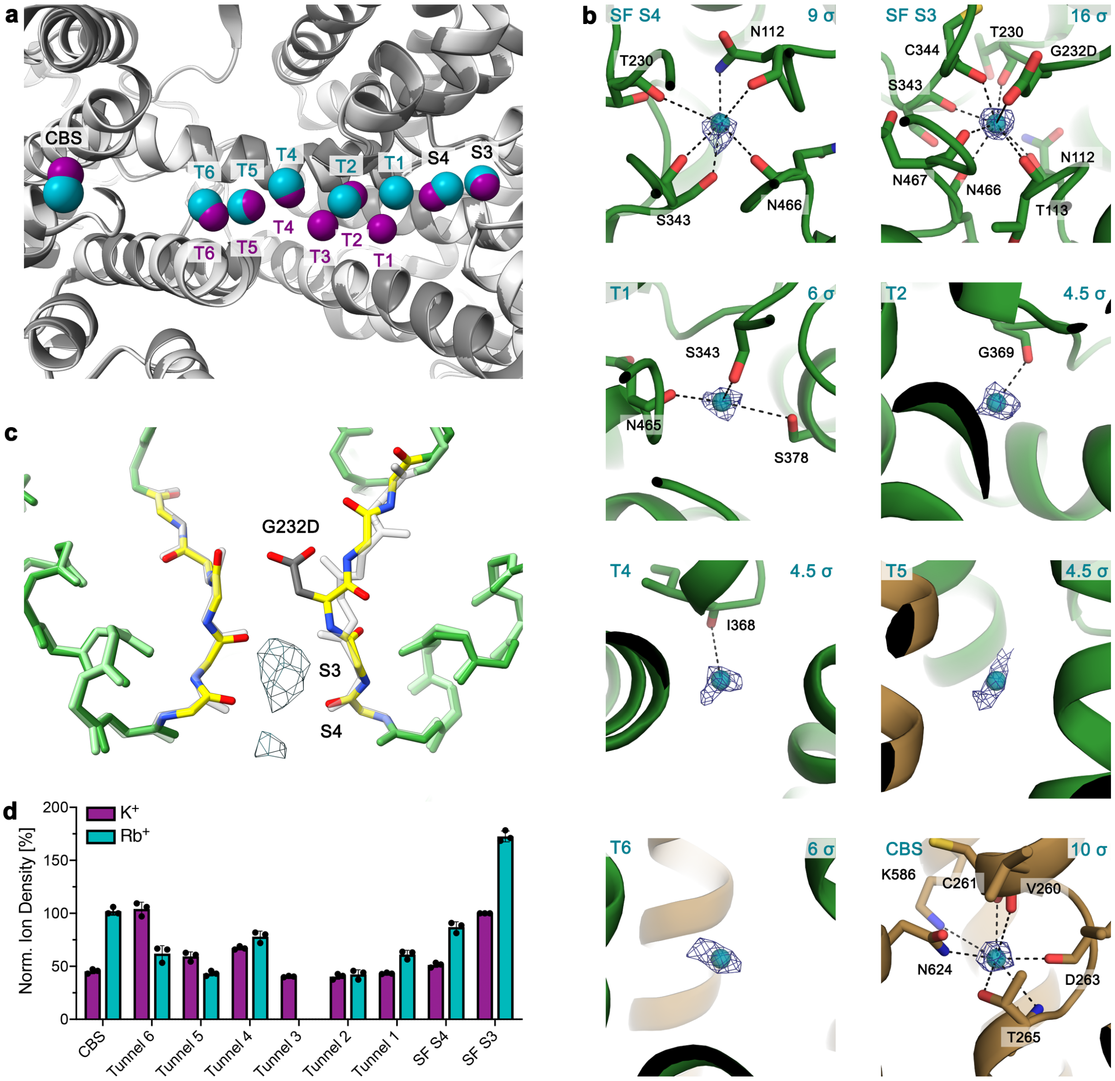

**Supplementary Figure 9: Ion coordination in Rb$^+$-loaded KdpFA$_{G232D}$B$_{S162A}$C. a,** Comparison of ion positions in Rb$^+$- and K$^+$-loaded E1·ATP structures of KdpFABC. Rb$^+$ ions shown in turquoise, K$^+$ ions in purple. Well-coordinated positions at the beginning and end of the intersubunit tunnel show a high correlation between the two structures, while positions between these optima with fewer coordinating residues show a higher deviation. Tunnel ion T3 was only identified in the K$^+$-loaded structure. **b**, Coordination of Rb$^+$ ions in the intersubunit tunnel. No coordinating residues were identified for Tunnel ions T5 and T6. Cryo-EM densities (mesh) shown with contour levels indicated in top right corner of each panel. **c,** SF of KdpFA$_{G232D}$B$_{S162A}$C shown in color, with wild-type SF from KdpB$_{D307N}$ underlaid in grey. The side chain of KdpA$_{G232D}$ inserts into SF coordination sites S1 and S2, blocking ion binding at these positions and possibly explaining the lowered affinity of KdpFA$_{G232D}$BC for ion substrates[8,9]. At the same time, the side chain adds a new coordination moiety to coordination site S3, changing the coordination geometry and explaining the reduced selectivity of the variant permitting Rb$^+$ passage through the SF. Ion densities (mesh) were observed in the S3 and S4 sites in the Rb$^+$-loaded structure of KdpFA$_{G232D}$B$_{S162A}$C. **d,** Density comparison of Rb$^+$ and K$^+$ positions identified in Rb$^+$-loaded KdpFABC and [EMD-12478], normalized to the K$^+$ density in SF S3. Error bars indicate the standard deviation from comparisons under three different map filter conditions.

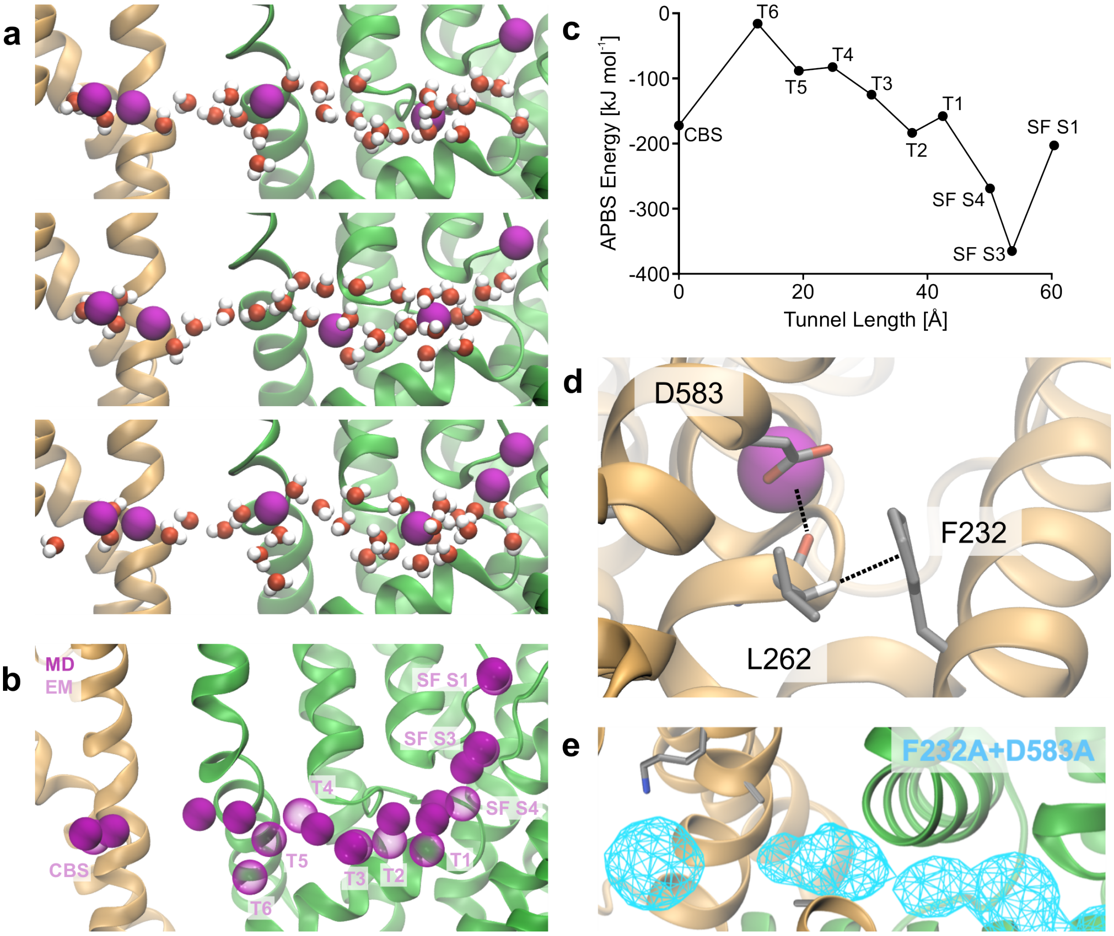

**Supplementary Figure 10: Analysis of ion occupancy in the intersubunit tunnel by MD simulations. a,** Snapshots of MD simulations at arbitrary time points showing unstructured waters aiding $K^+$ coordination in the intersubunit tunnel. **b,** Overlay of all positions used for APBS analysis of MD-relaxed ion positions (solid purple), compiled from multiple snapshots. The input ion coordinates from cryo-EM are shown in transparent purple. **c,** Coordination energy of ion positions from cryo-EM without MD relaxation by Adaptive Poisson-Boltzmann Solver (APBS) analysis. All positions observed in the cryo-EM structure are energetically favorable, although positions T4-T6 are less so, and are immediately abandoned in MD simulations. **d,** Coordination near $KdpB_{D583}$ aided by the backbone carbonyl of $KdpB_{L262}$, which is in turn CH-π stacked with $KdpB_{F232}$. **e,** Ion progression through intersubunit tunnel in $KdpFAB_{F232A/D583A}C$. Removal of the steric hindrance of the phenylalanine allows ion passage of $K^+$ even in the absence of the $KdpB_{D583}$ energy well.

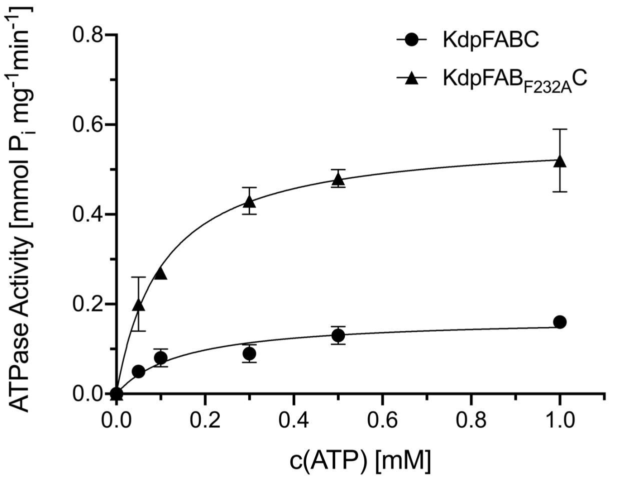

**Supplementary Figure 11: Michaelis-Menten kinetics of ATP hydrolysis by KdpFABC and $KdpFAB_{F232A}C$.** Michaelis-Menten kinetics of KdpFABC ATP turnover, showing no effect of the mutation $KdpB_{F232A}$ on the apparent affinity for ATP ($K_{m,ATP}$ 0.15 mM for KdpFABC, 0.10 mM for $KdpFAB_{F232A}C$) but a threefold increase in the $V_{max}$ (0.17 mmol $P_i$ $mg^{-1}min^{-1}$ for KdpFABC, 0.58 mmol $P_i$ $mg^{-1}min^{-1}$ for $KdpFAB_{F232A}C$). This demonstrates that the observed increase in ATPase rate is not caused by an increased affinity for ATP. Error bars indicate the standard deviation of technical triplicate measurements.

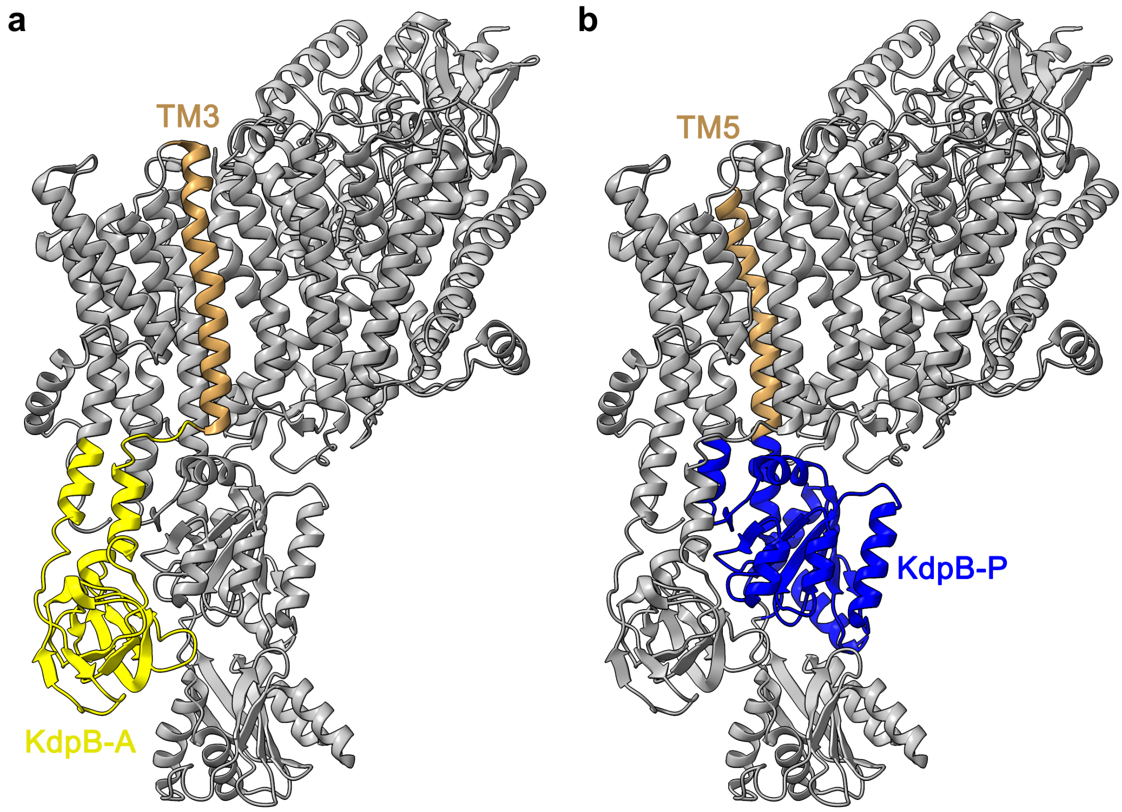

**Supplementary Figure 12: Connection of helices containing residues involved in ATPase coupling to the cytosolic domains of KdpB.** KdpFABC in the ribbon conformation, with relevant TM helices and cytosolic domains of KdpB highlighted. KdpF is excluded for visibility. **a,** KdpB TM3, which harbors $KdpB_{F232}$, is connected to the A domain, providing the structural basis for the proposed regulation of A domain plasticity by this residue. **b,** KdpB TM5 contains $KdpB_{D583/K586}$, which are proposed to be responsible for stimulation of ATP hydrolysis, and is connected to the P domain, which enacts ATP hydrolysis. Conformational rearrangements of these residues can thus be allosterically transferred to the P domain to initiate autophosphorylation.

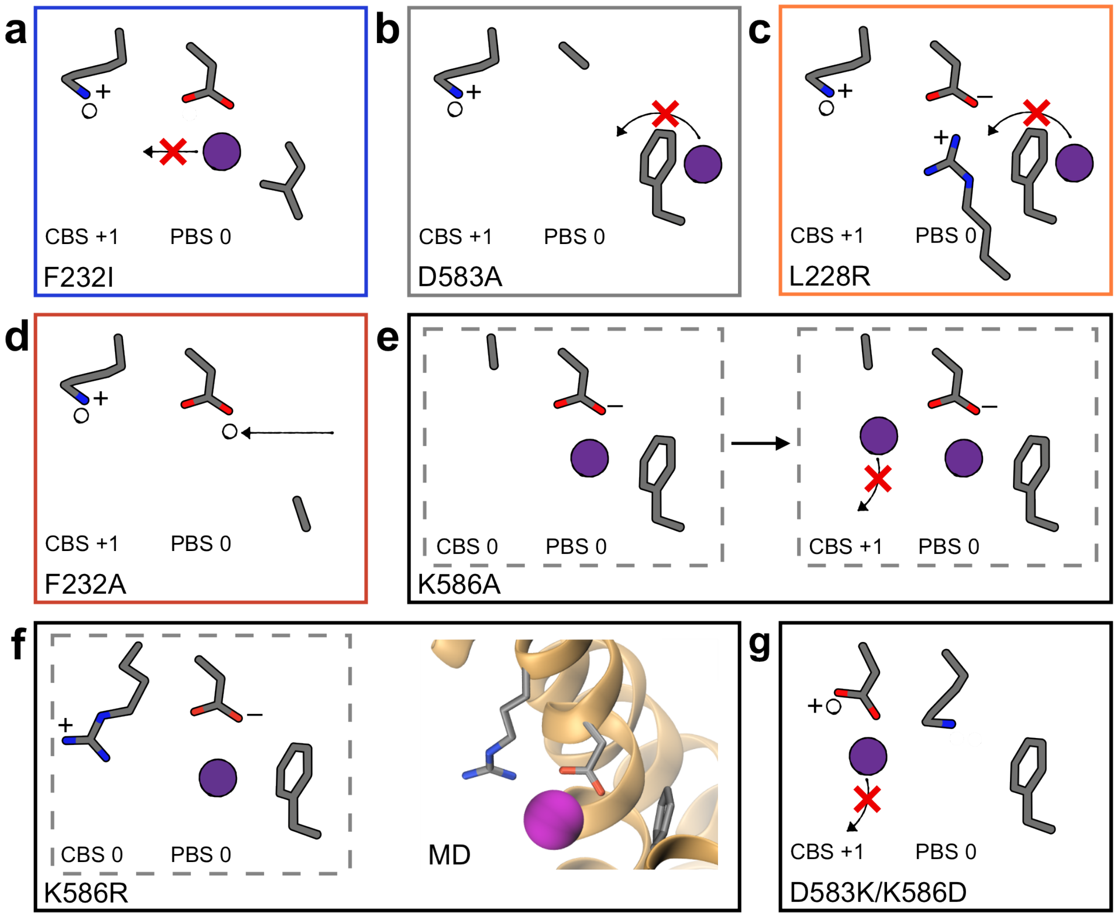

**Supplementary Figure 13: Explanation of mutation phenotypes in the context of our mechanistic model. a,** KdpB$_{F232I}$ shows wild-type levels of ATP hydrolysis and K$^+$-dependence, but a slower transport rate. This is due to the loss of cation-π interactions with the phenylalanine side chain, preventing ion forwarding to the CBS, whilst the isoleucine side chain still constitutes a steric hurdle in the tunnel. **b**, KdpB$_{D583A}$ has constitutive ATPase activity, independent of K$^+$ and with an increased resistance to orthovanadate, and abolishes transport[23]. In this variant, the net charge in the CBS is +1, leading to constant ATPase activity, even without K$^+$. Furthermore, the charge at the PBS is lost, meaning K$^+$ is not pulled past KdpB$_{F232}$, explaining the lack of transport. The E2/E1 transition is accelerated, possibly because the displacement of ions from the CBS in the E2 state no longer needs to take place. **c,** Like KdpB$_{D583A}$, KdpB$_{L228R}$ causes orthovanadate-resistant ATP activity uncoupled from K$^+$, no transport, and an accelerated E2/E1 transition. This is likewise due to a neutralization of the PBS. Ion progression from the intersubunit tunnel is prevented by neutralization of the energy well at the PBS, explaining the lack of transport. No ions reach the CBS, likely accelerating the E2/E1 transition. **d,** KdpB$_{F232A}$ leads to an increased ATPase rate independent of K$^+$ and a slowed transport rate. The loss of the phenylalanine side chain removes the gatekeeper for the PBS, allowing unspecific protonation of the aspartate from unspecified donors in the intersubunit tunnel. This leads to a neutralization of charges in the PBS, stimulating ATP hydrolysis in the absence of K$^+$. Protonation of KdpB$_{D583}$ also neutralizes the energy well, decelerating ion progression towards the CBS, in part explaining the slower transport rate. Additionally, cation-π interactions required for ion forwarding are lost, also contributing to the lowered transport rate. The increase in ATPase activity may be due to deregulation of the A domain by the loss of the sterically demanding side chain in the intersubunit tunnel. **e,** KdpB$_{K586A}$ significantly reduces the ATPase activity of the complex, although the remaining activity is still dependent on K$^+$[83]. No transport was observed. A single ion entering the PBS in this variant gives a net charge of 0, meaning there is no ATPase stimulation. However, this variant in principle could allow two ions to pass KdpB$_{F232}$, with one ion binding at the CBS and one at the PBS. This is expected to be a rare occurrence explaining the low level of K$^+$-stimulated ATP hydrolysis. However, displacement of the CBS ion in the E2 state is impossible due to the loss of the KdpB$_{K586}$ side chain, resulting in the lack of transport. **f,** KdpB$_{K586R}$ shows wild-type levels of ATP hydrolysis, K$^+$-

dependence, and transport[23,83]. While a protonation inversion with the arginine side chain is less likely than with lysine, the arginine is able to aid coordination of the CBS ion with the face of its delocalized electron system (left panel), and is long enough to potentially form a salt bridge with $KdpB_{D583}$ instead of switching protonation states. This is supported by MD simulations (right panel). **g**, Most puzzling are the results from the inversion of the CBS dipole ($KdpB_{D583K/K586D}$), which resulted in normal ATPase activity but abolished transport[83]. It is possible that the double mutation causes significant rearrangements in the CBS, changing the behavior of ions in this section of the complex. If a $K^+$ ion somehow reaches into the CBS, possibly pulled by the negatively charged $KdpB_{K586D}$, a subsequent protonation switch would establish the charge distribution required for ATP hydrolysis. At the same time, the lysine side chain required for ion displacement is lost, explaining the lack of transport by this variant.

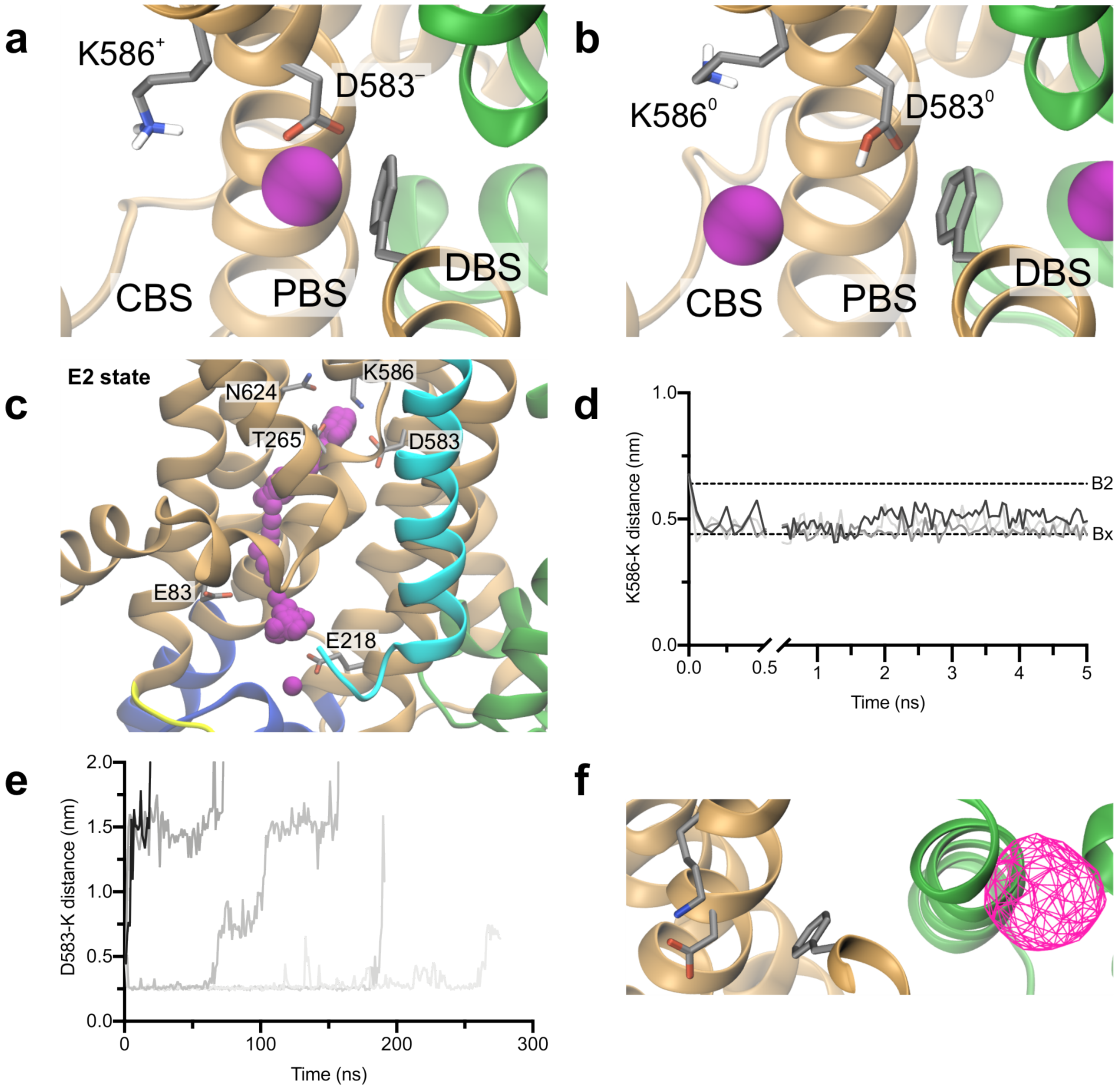

**Supplementary Figure 14: Progression of ions between different steps of the transport cycle in MD simulations. a** and **b,** Simulations of ion behavior in different charge states of the CBS in the E1·ATP state. **a,** Snapshots of the DBS, PBS and CBS in the E1 ground state, with both $KdpB_{D583}$ and $KdpB_{K586}$ charged. The ion is stably bound by $KdpB_{D583}$, supported by $KdpB_{F232}$. **b,** When protonation between $KdpB_{D583/K586}$ is inverted (both neutral) and a second ion approaches the other face of $KdpB_{F232}$, the first ion is repelled forward and stably occupies the CBS. **c, d,** and **e,** Simulations of ion behavior in the CBS and intersubunit tunnel in the E2 conformation. Simulations were run using the 2.9 Å resolution $E2{\cdot}P_i$ structure [7BGY]. **c,** Sequential ion positions showing the smoothed ion release pathway from the CBS in a single simulation of the E2 state. In this conformation, ions in the CBS are released from the complex through an inward-open half-channel. Ions placed in the CBS exit the complex without entering the previously proposed low-affinity release site adjacent to the CBS. Ions placed in this site rapidly move back to the CBS before following the same exit route (data not shown). **d,** Timescales of $K^+$ occupancy of the low-affinity (B2) site in the E2 state from three simulations. Ions rapidly leave the B2 site towards the first observed CBS site (Bx), where occupancy is more stable. **e,** Timescales of ion release from the CBS in the E2 state from five simulations. Ion displacement from the CBS can take up to 250 ns or longer. **f,** Ion progression through the intersubunit tunnel in the E2 state. Alternating access is facilitated by a constriction at the KdpA/KdpB interface, which prevents ions from the intersubunit tunnel from progressing towards the CBS.

```
P03960 - E. coli                  204 IAMVEGAQRRKTPNEIALTILLIALTIVFLLATATLWPFSAWGGNA 249
Q7NN40 - G. violaceus             219 IALIEGAKRQKTPNEIALTVLLAVLTLIFLIVVATLPPIAAFVGAP 264
B0R9M0 - H. salinarum             209 IGLVEDAQRQKTPNEIAMTILLSGLTLVFVVAVATMFFFGEYLASF 254
A0A0H3C7Y6 - C. vibrioides        211 IAMVEGADRRKTPNEIALAVLLAGLTLIFLIAVVTLLGPGKFSGVA 256
P63682 - M. bovis                 221 IALVEGAARQQTPNEIALNILLAGLTIIFLLAVVTLQPFAIYSGGG 266
P57699 - H. salinarum             209 IGLVEDAQRQKTPNEIAMTILLSGLTLVFVVAVATMFFFGEYLASF 254
B0JJ96 - M. aeruginosa            229 ISLVEGAERTKTPNEIALTVLLAVLTQVFLIVVATIPPIGNYIAGF 274
Q6GKN3 - S. aureus                201 IGLVEGATRKKTPNEIALFTLLMTLTIIFLVVILTMYPLAKFLNFN 246
Q725T7 - D. vulgaris              199 IALVEGAERKKTPNEIALNILLAGLTLIFILATVTLKPMALFHGAR 244
Q8YPE9 - Nostoc sp.               226 IALVEGAERSKTPNEVALTVLLAVLSLVFLFVIATLPAFAYYADTP 271
Q02CX6 - S. usitatus              220 IALVEGAQRQKTPNEIALNIVIAGLTLVFLLAVVTLQPFAIYSVAT 265
P73867 - S. usitatus              218 IDLVEGAERSKTPNEIALTVLLAVLTLVFLIVVATLPPPANYIDSP 263
C8WRA9 - A. acidocaldarius         205 IALVEGASRQKTPNEIALSVLLAGLTLIFLIVIDCLPPIAKGLGAH 250
C1FA48 - A. capsulatum             205 IALVEGTQRQKTPNEIALNILLAGLTIIFLLATVTLQPFAIYSGAP 250
B1MDL0 - M. abscessus              203 IALVEGASRQKTPNEIALNILLASLTIIFLLAVVALGPMGNYGGEQ 248
B7JY04 - R. orientalis             228 IALVEGAERTKTPNEIALTVLLAVLTQVFLVVVATIPPIAHYVGSP 273
B5EH79 - G. bemidjiensis           207 ISLIEGAKRRKTPNEIALEVLLIALTLVFLLVCANISPLSVYSVKA 256
Q9R6X1 - Anabaena sp.             226 IALVEGAERTKTPNEVALTVLLAVLSLVFLFVVATLPVFAYYADTP 271
Q8A520 - B. thetaiotaomicron      201 IALVEGASRQKTPNEIALTILLAGFTLVFVIVCVTLKPFADYSNTV 246
Q97BF6 - T. volcanium             195 IELVEKSTREKTPNEISLTVFLSGLTLIFLVITASIFAISHYFGRT 240
Q5KUV4 - G. kaustophilus          202 ISLVEGATRQKTPNELALNILLVTLTLIFLIVVVTLVPIARYVGIH 247
Q1IUD4 - K. versatilis            198 IALVEGAERQKTPNEIALNILLAGLTIIFLLAVVTLQPFAIYSGAQ 243
Q74AA9 - G. sulfurreducens        207 ISMIEGAKRRKTPNEIALEVLLIALTAVFLLVCANISPLSVYSVRA 252
```

**Supplementary Figure 15: Sequence alignment of KdpB sequences around KdpB$_{F232}$.** KdpB sequences from different species, listed by UNIPROT accession number and species, were aligned using Clustal Omega[81] and colored by degree of conservation compared to *E. coli* KdpB (dark green – highly conserved; red – not conserved). Species were chosen to reflect the range of genetic diversity, including gram-negative and gram-positive bacteria[15]. The highly conserved KdpB$_{F232}$ is indicated by the pink marker.

**Supplementary Table 1: Cryo-EM data collection, refinement and validation statistics**

| | KdpFAB$_{D307N}$C<br>E1·ATP, $K^+$<br>(EMD-12478)<br>(PDB 7NNL) | KdpFA$_{G232D}$B$_{S162A}$C<br>E1·ATP, $Rb^+$<br>(EMDB-12482)<br>(PDB 7NNP) |
|---|---|---|
| **Data collection and processing** | | |
| Magnification | 49,407 | 49,407 |
| Voltage (keV) | 200 | 200 |
| Electron exposure ($e^-/Å^2$) | 52 | 52 |
| Defocus range (μm) | -0.5 to -2.0 | -0.5 to -2.0 |
| Pixel size (Å) | 1.012 | 1.012 |
| Symmetry imposed | C1 | C1 |
| Initial particle images (no.) | 331,673 | 756,834 |
| Final particle images (no.) | 160,776 | 196,682 |
| Map resolution (Å) | 3.1 | 3.2 |
| FSC threshold | 0.143 | 0.143 |
| Map resolution range (Å) | 2.9-4.7 | 3.1-5 |
| **Refinement** | | |
| Initial model used (PDB code) | 6HRA | 6HRA |
| Model resolution (Å) | 3.3 | 3.4 |
| FSC threshold | 0.5 | 0.5 |
| Model resolution range (Å) | 80-3.1 | 80-3.2 |
| Sharpening B-factor ($Å^2$) | -92 | -130 |
| Model composition | | |
| Non-hydrogen atoms | 11099 | 11096 |
| Protein residues | 1456 | 1456 |
| Ligands | K: 10 | Rb: 8 |
| | CDL: 2 | CDL: 2 |
| | ACP: 1 | ACP: 1 |
| B factors ($Å^2$) | | |
| Protein | 63.9 | 47.0 |
| Ligand | 65.2 | 48.3 |
| R.m.s. deviations | | |
| Bond lengths (Å) | 0.008 | 0.007 |
| Bond angles (°) | 0.885 | 0.922 |
| Validation | | |
| MolProbity Score | 1.84 | 1.77 |
| Clash score | 10.82 | 11.08 |
| Poor rotamers, % | 0.35 | 0.00 |
| Ramachandran plot | | |
| Favored (%) | 95.78 | 96.69 |
| Allowed (%) | 4.22 | 3.31 |
| Outliers (%) | 0.00 | 0.00 |
